# A Novel Prognosis Model based on Comprehensive Analysis of Pyroptosis-Related Genes in Breast Cancer

**DOI:** 10.1101/2022.04.05.486932

**Authors:** Shijie Zeng, Yuefei Wang, Yixi Yang

## Abstract

**Background:** Pyroptosis plays a dual role in cancer. It can not only induce chronic tumor necrosis, but also stimulate acute inflammation to enhance immune response. However, the study of cell pyroptosis in Breast Cancer (BC) is still limited.

**Methods:** A total of 26 pyroptosis-related differential genes were obtained, and PPI network and correlation analysis were demonstrated as well. Besides, 3 pyroptosis-related prognostic genes were collected by the univariate COX analysis. Through these 3 genes, patients were clustered into 2 sub-types by the Consensus clustering, then the EM and XMeans were also carried out to recheck the result. Furthermore, the significant prognosis-related genes were obtained by the Random Forest and LASSO analysis. Based on the above, the corresponding weights were calculated through the univariate and multivariate Cox analysis, and the prognostic model was constructed. In addition, the ROC curve, risk curve, PCA, t-SNE, COX, GO, KEGG and ssGSEA were analyzed, respectively.

**Results:** Through the verification of 3 different clustering algorithms, 2 sub-types of BC were obtained. Furthermore, ‘ELOVL2’, ‘IGLV6-57’, ‘FGBP1’, ‘HLA-DPB2’ with weights of −0.183, −0.101, −0.227 and −0.254 were employed to establish the prognostic model. Validation shows that our model has good effect and the Go KEGG and Immune microenvironment analysis showed the enrichment of the differential genes.

**Conclusion:** Our study identified 4 genes that are closely correlated with the overall survival of BC patients, and the prognostic model can effectively divide patients into high- and low-risk groups, which may have certain guiding significance for prognosis.

## Introduction

As of 2020, Breast Cancer (BC) has caused 685 thousand deaths and is expected to reach 4 million 400 thousand cases by 2070^[1, 2]^. Today, BC remains one of the major diseases facing the world. Moreover, there are significant differences in morbidity and mortality between countries. For example, in terms of incidence, the lowest such as Iran, 35.8 cases per 100,000 people; while the highest, such as Belgium, with 112.3 cases per 100,000^[3]^. In short, BC has become one of the major diseases threatening women.

In recent years, the survival rate of patients has been effectively improved by means of chemotherapy, endocrine therapy, targeted drug therapy and radiotherapy. However, the prognosis of patients remains to be improved ^[4–6]^. Therefore, it is urgent to establish a model to predict and evaluate the risk and treatment effect of BC patients.

Pyroptosis is a recently discovered mode of programmed cell death, which is characterized by the continuous expansion of cells until the rupture of cell membrane, resulting in the release of cellular contents that activate intense inflammatory responses ^[7, 8]^. As the trigger mechanism of pyroptosis, inflammasome activation is mainly induced by the canonical caspase-1 inflammasome pathway and the non-canonical caspase 4/5/11 inflammasome pathway ^[9, 10]^. More and more studies have shown that similar programmed cell death (such as pyroptosis, apoptosis, necroptosis, etc.) is closely related to the treatment of cancer.

In this study, by using TCGA (Texas Cotton Ginners’ Association) and GEO (Gene Expression Omnibus) cohort, BC patients were divided into two sub-types based on 3 *pyroptosis-related prognostic genes*, and there were significant differences in survival between these two sub-types. In order to further analyze this survival situation, a prognostic model was established to divide the patients into high- and low risk groups for prognosis. Then, the effect of the prognostic model was analyzed by ROC curve, survival curve, risk curve, PCA analysis, and univariate and multivariate Cox analysis. Finally, the differences in biological function and immune microenvironment between high and low-risk patients were explored.

## Method and Materials

### Collection of Data

The flowchart of this paper is shown in the Figure 1. The complete RNA sequencing (RNA-seq) and clinical characteristics of patients were obtained from 113 normal cases and 1109 BC patients from the TCGA-BRCA project (https://portal.gdc.cancer.gov/repository) on 9 November 2021. The data of the external validation cohort were downloaded from the GEO database, which include RNA-seq data and clinical data of 104 BC and 17 normal breast biopsies. In addition, the maximal follow-up was 3,026 days, and minimal follow-up was 138 days with a mean follow-up of 1,887 days. (https://www.ncbi.nlm.nih.gov/geo/, ID: GSE42568).

**Figure 1.**
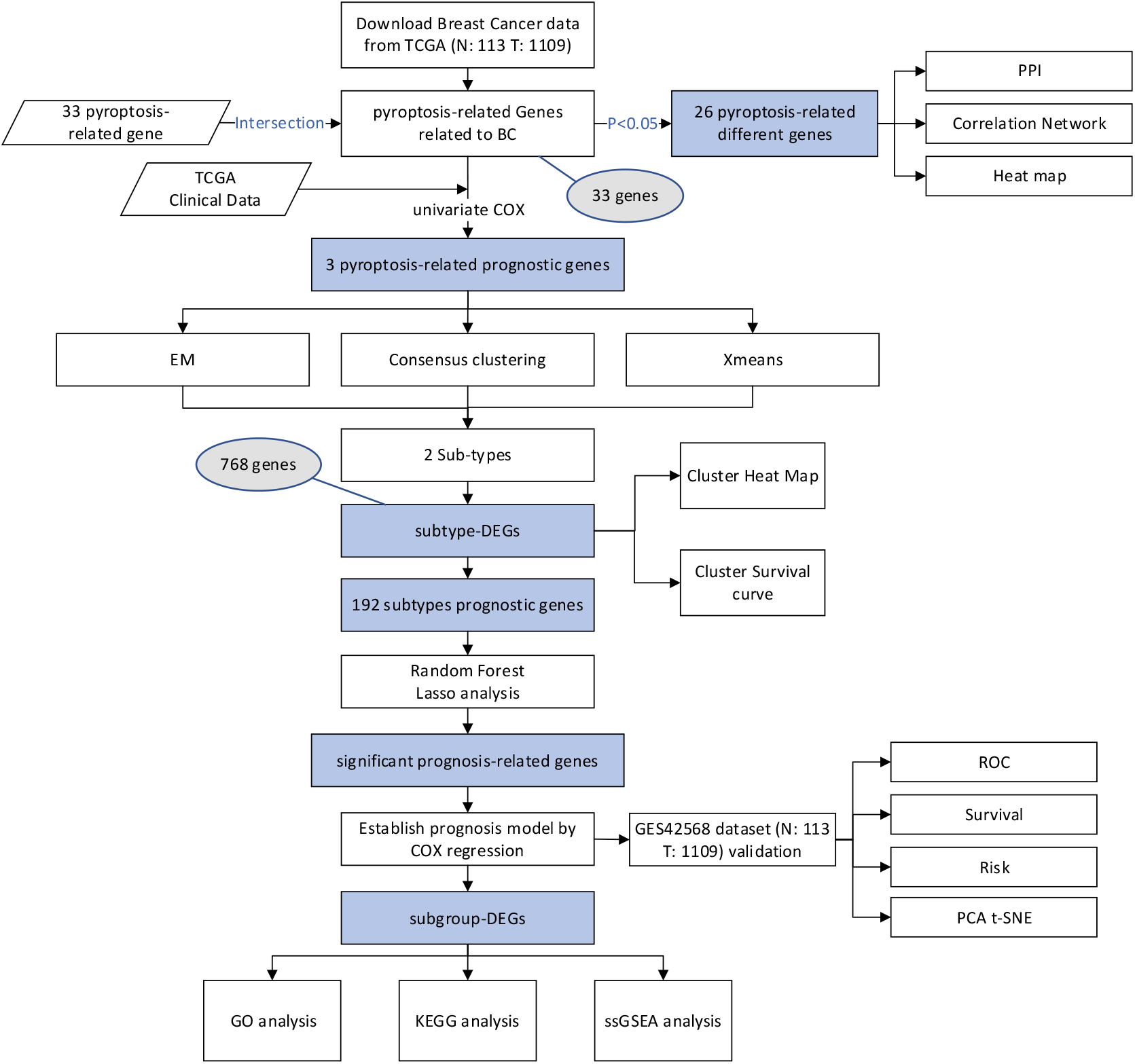
The flowchart of this research.

### Pyroptosis-Related Genes

According to the previous literatures, a total of 33 *pyroptosis-related genes* were collected, and they were displayed in detail in the Table 1. Then, the names of 33 *pyroptosis-related genes* were compared with those in the Gene Expression Matrix which transformed from the complete RNA sequencing (RNA-seq) data, and verified that these genes are related to BC disease through the intersection operation.

**Table 1.**
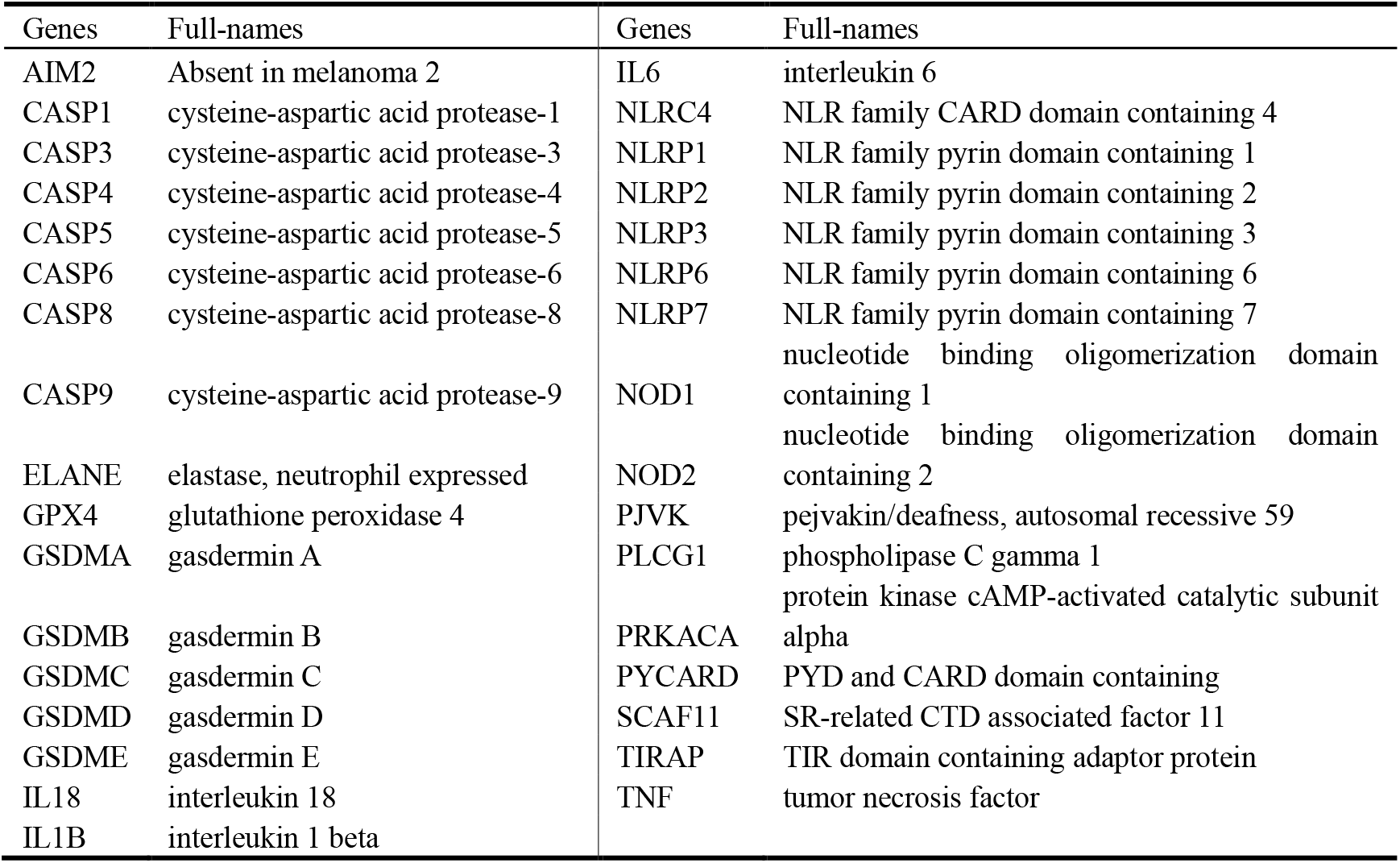
Pyroptosis-related genes.

### Identification of DEGs (Differentially Expressed Genes)

The DEGs were identified through the “limma” package of R language, and there are different types of DEGs according to different requirements in the whole process. For example, DEGs between normal tissue and breast tumor tissue, different sub-types, and different risk groups. These DEGs were confirmed by the Mann–Whitney–Wilcoxon Test method, which threshold was set to *p* < 0.05 and |logFC| < 1. Moreover, the results were demonstrated by heat maps, and the degree of difference was marked with the label *, where * denotes *p* < 0.05, ** denotes *p* < 0.01, and *** means *p* < 0.001.

### Protein-Protein Interaction (PPI) and Correlation Network analysis

To further explore the interactions of these *pyroptosis-related genes* in BC, the PPI network for the DEGs was constructed with Search Tool for the Retrieval of Interacting Genes (STRING), version 11.0 (https://string-db.org/). Besides, the threshold value was set at 0.9 (the highest confidence). In addition, Correlation Network analysis with a threshold of 0.2 was carried out to analyze the correlation between genes and explore the differences between positive and negative correlations.

### Univariate and multivariate Cox regression analyses

The Significant DEGs were identified from the prognosis-related DEGs by the univariate Cox regression analysis. In addition, both univariate and multivariate analyses were used for establishing the prognostic model. The Hazard Ratio, 95% confidence interval, and p-values were determined separately.

### Unsupervised Clustering Analysis

Clustering is a type of unsupervised technology, through which data can be divided into multiple sub-categories. In this paper, the Consensus Clustering was performed to classify the 1089 BC patients in the Gene Expression Matrix into two sub-types by 3 *pyroptosis-related prognostic genes*, and there were 398 and 691 patients with two subtypes, respectively. In addition, the EM and XMeans method were used to verify the accuracy of the clustering. The results showed that it was more appropriate to divide the patients into two subtypes.

### Identification of prognosis-related DEGs

In order to better explore the risk and survival of patients, the Random Forest and Lasso Regression analyses were used to screen prognosis-related DEGs. The Random Forest with 1000 trees was established by the “vh” method to extract variables with large variances in the analysis results. Then, the Lasso Regression was performed with 10-fold cross-validation and the genes that regression coefficients are not equal to 0 were extracted. Finally, the Radom Forest extracting genes and the lasso regression extracting genes were intersected as the significant prognostic genes, which num is 4.

### Establishment and validation of the prognostic model

The Univariate and Multivariate Cox regression analyses were also used to establish the prognostic model of 4 *significant prognosis-related genes*. The univariate Cox analyses found that the *p* value of all these genes was less than 0.05. Then, the multivariate Cox regression was used to calculate their coefficients for the prognostic model. The risk score in the prognostic model equation is as follows:

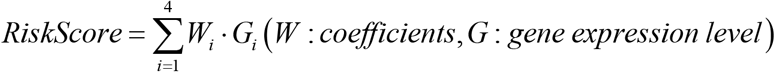

According to the above formula, the TCGA and GEO BC patients can be divided into low- and high-risk sub-groups by the median value of risk scores.

### The Survival, ROC, Risk, and PCA analysis

Based on different risk groups, some validation work was done to test the accuracy of the proposed prognostic model. First, the overall survival of high and low-risk groups was compared by the Kaplan-Meier curves and log-rank tests. Then, the AUC values of ROC in 3-, 5- and 7-years were calculated to validate the accuracy of this model. Next, the PCA was conducted to overview the different distribution patterns between two sub-groups. Moreover, the distribution of patients from different risk groups was verified, and the effect of the model was confirmed through the division of the curve. Finally, the univariate and multivariate Cox regression were used to verify if the model can be used as the prognostic factor.

### Functional enrichment analysis of DEGs between low- and high-score groups

To explore the potential biological functions and pathways among the high- and low-risk sub-groups, the DEGs were filtered according to specific criteria (|log2FC| ≥ 1 and FDR< 0.05). Based on these DEGs, GO and KEGG analyses were conducted with the “clusterProfiler” package.

### Immune microenvironment analysis

To investigate the molecular feature among the high- and low-risk groups, the Gene Set Variation Analysis (GSVA) enrichment analysis was established by using the “GSVA” R packages.

### Statistical analyses

All visualization and statistical analyses were performed using the R (R 4.1.1), and some clustering analyses were conducted by the weka (weka 3.8.5) software. Using the Wilcoxon Test to extract the DEGs under the three scenarios of different sub-types of BC, different sub-groups of high- and low-risks, and normal tissue and breast tumor tissue. Independent prognostic factors were identified by the univariate and multivariate Cox proportional hazards regression analyses. The Kaplan–Meier method was used to compare the overall survival between low- and high-risk sub-groups. The *p* < 0.05 was regarded to be significant.

## Results

### Identification of different pyroptosis-related genes in normal and BC tissues

The gene expression matrix from TCGA cohort was intersected with 33 *pyroptosis-related genes* by gene names. With the intersected Gene Expression Matrix, the Mann–Whitney–Wilcoxon Test method was performed by setting |log2FC| > 1 and FDR<0.05 as the cutoff value to select the *pyroptosis-related differential genes* in normal and BC tissues, and the corresponding heatmap was plotted (Figure 2A). Meanwhile, the PPI network was constructed by the STRING. The minimum required interaction score was set to 0.9 (the highest confidence), and CASP1, PYCARD, NLRC3 were identified as hub genes (Figure 2B). The correlation network consisting of 26 *pyroptosis-related differential genes* was presented in Figure 2C (correlation coefficient > 0.2, positive correlation was shown with red line, negative correlation was shown with blue line). In addition, in order to explore the *pyroptosis-related prognostic genes* in patients, the univariate Cox regression was conducted to obtain the three genes of CASP9, IL18, TIRAP, these were recognized as the genes that cause differences between patients.

**Figure 2.**
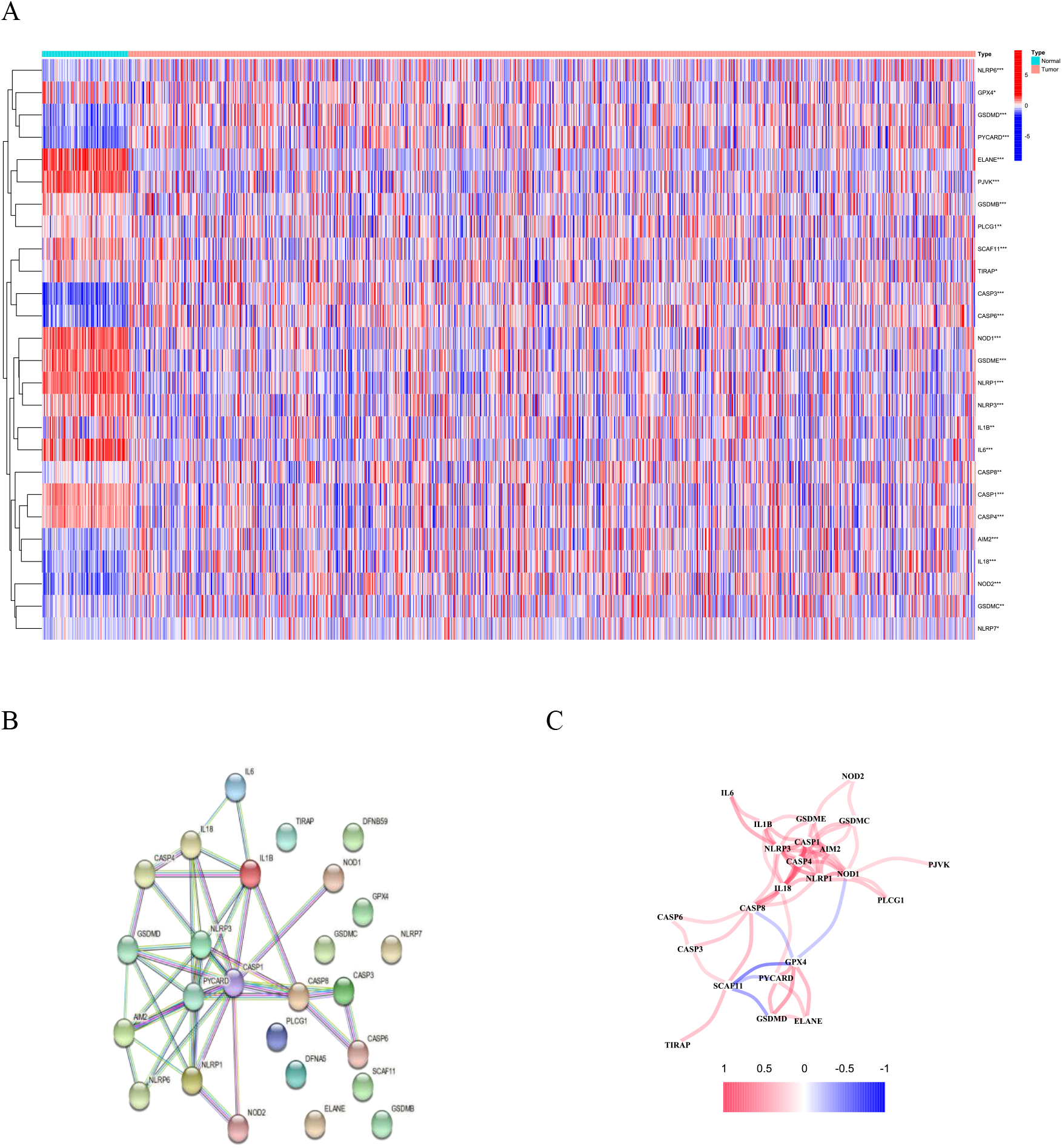
Expression and interaction of pyroptosis-related genes. **A** The heatmap of cell pyroptosis-related genes between normal tissue (N, blue) and tumor tissue (T, red) (Blue indicates low expression level, while red indicates high expression level). **B** PPI network, Interaction of pyroptosis-related genes (interaction score = 0.9). **C** The correlation network of pyroptosis-related genes (Red line indicates positive correlation, while blue line indicates negative correlation. Color depth reflects correlation strength)

### Tumor clustering based on DEGs

Clustering was used for identifying the subtypes of BC. In this study, different clustering methods were conducted to explore the number of BC sub-types and corresponding patients. Firstly, the Consensus Clustering were performed on 1089 BC patients from the TCGA cohort. The algorithm does not specify the number of clusters, but verifies the clustering effect within the assigned range. Since 3 *pyroptosis-related prognostic genes* of ***CASP9*, *IL18*** and ***TIRAP*** were significantly correlated with the prognosis of BC, the values of the above 3 genes of the patients were input to the Consensus Clustering. By setting the interval of *k* to [2,9], it was found that when the parameter is equal to 2, the intragroup correlation is relatively high, while the inter-group correlation is quite low, which means that it is more appropriate to set 2 as the number of BC subtypes (Fig. 3a). In addition, EM and XMeans algorithms were performed to verify the results of clustering, these two algorithms also do not need to specify the number of subtypes, and automatically summarize the data into different categories by iteration. (Table 2). Moreover, the survival curves of different sub-types were demonstrated in Figure 3B. Besides, gene expression profile and clinical data with the attributes including Age (≤65 years or> 65 years old), Gender, Stage, T, M, N, fustatus, and Subtype, as shown in the heat map (Figure 3C).

**Figure 3.**
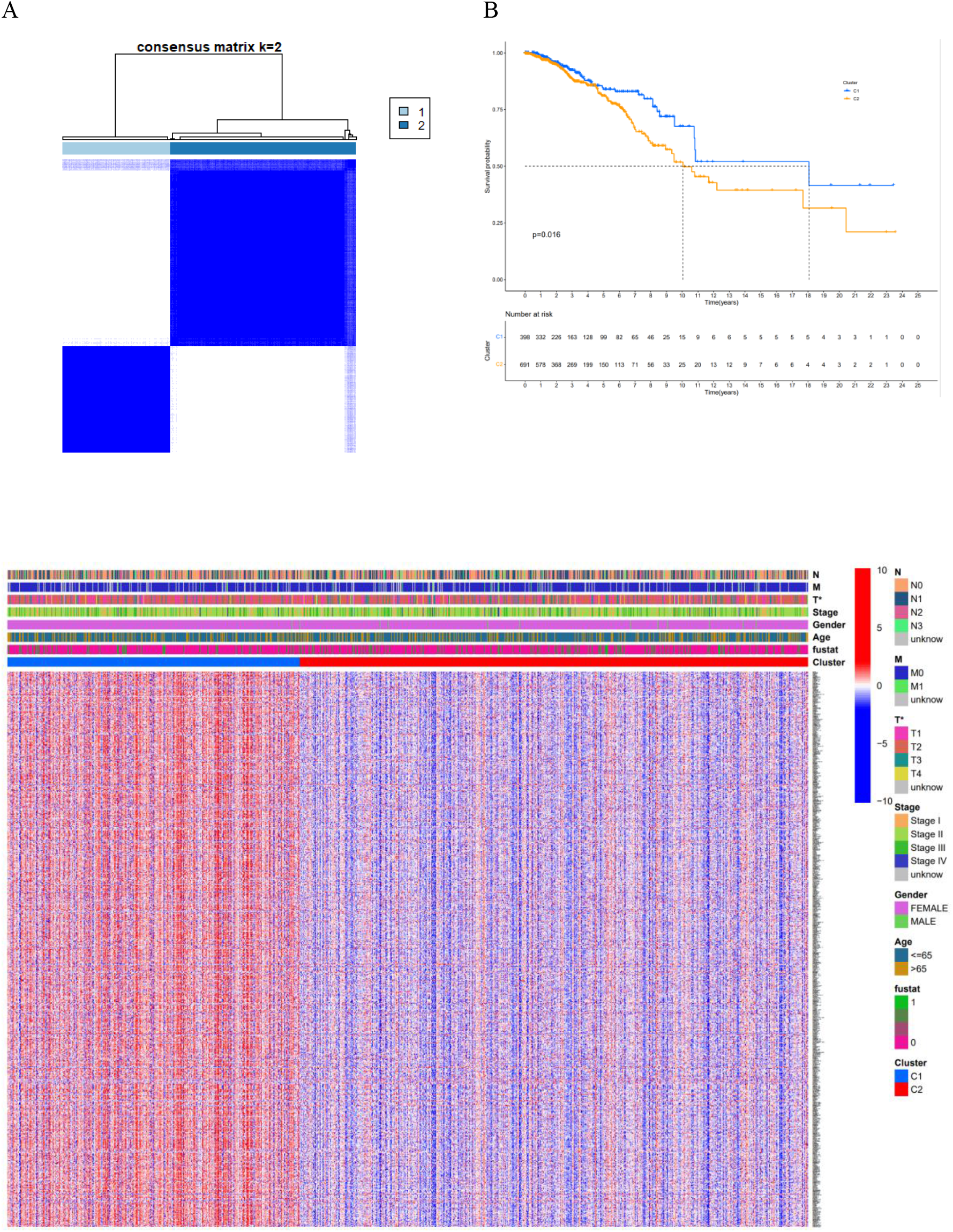
Tumor classification based on the pyroptosis-related DEGs. **A** 1085 BC patients were grouped into two clusters according to the Consensus Clustering matrix (maxK = 9). **B** Overall survival curves for the two clusters. **C** Heatmap and the clinical features of the two clusters classified by these DEGs (pvalue<0.05, “*”).

**Table 2.**
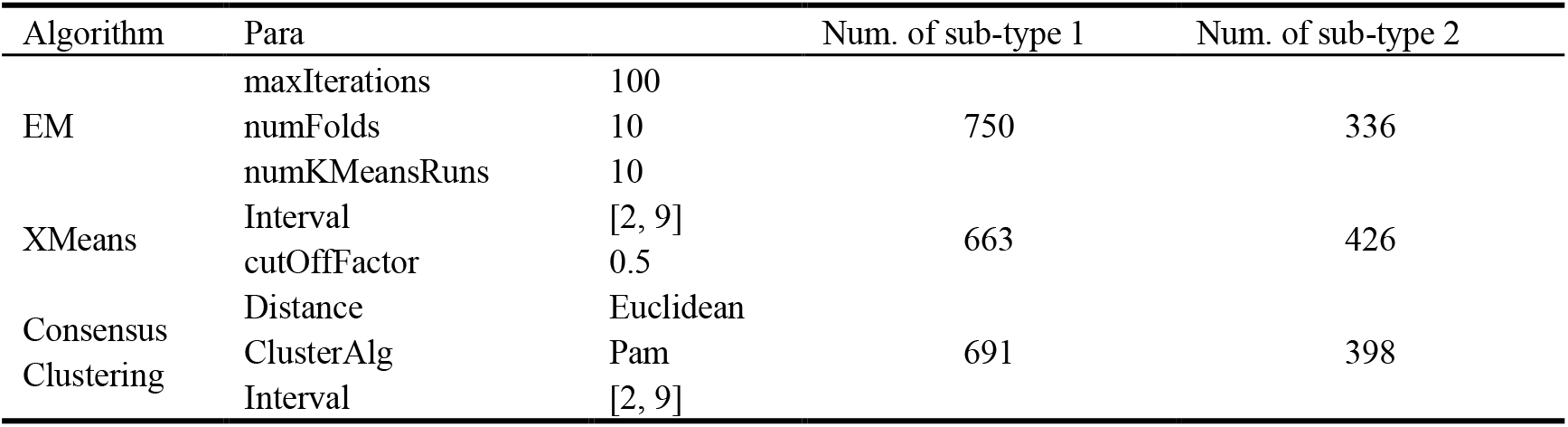
Parameters and results of three Clustering Algorithms

### Establishment of prognostic gene model

To analysis the survival differences of the 2 BC sub-types, the Mann–Whitney–Wilcoxon Test were adopted by setting |log2FC| > 1 and FDR < 0.05 as the cutoff value, and identified 768 *subtype-DEGs*. TCGA and GEO gene expression data were merged by normalizing and removing batch effects, and extracted the *merged-DEGs*, which combined from two cohorts and obtained 17577 genes. Furthermore, the *subtype-DEGs* was intersected with the *merged-DEGs* and finally got 500 *Same-Genes*, which can descript the differences between the 2 sub-types.

In the modeling process, first, the *Same-Genes* expression matrix was merged with corresponding survival time and survival status. According to the univariate Cox analysis, 192 *subtypes prognostic genes* were identified. Then, the *significant prognosis-related genes* were screened by the Random Forest and Lasso Regression, which selected 35 and 19 genes, respectively. Next, two sets were intersected and 4 final *significant prognosis-related genes* (Figure 4A), which were “ELOVL2”, “IGLV6-57”, “FGBP1” and “HLA-DPB2” were collected. Besides, the univariate Cox analyses and multivariate Cox regression were applied to calculate their coefficients for the prognostic model. The risk score of each patient can be calculated through the following formula:

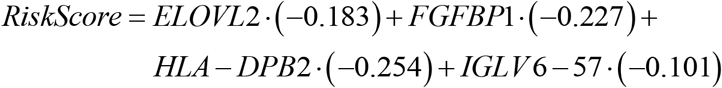

**Figure 4.**
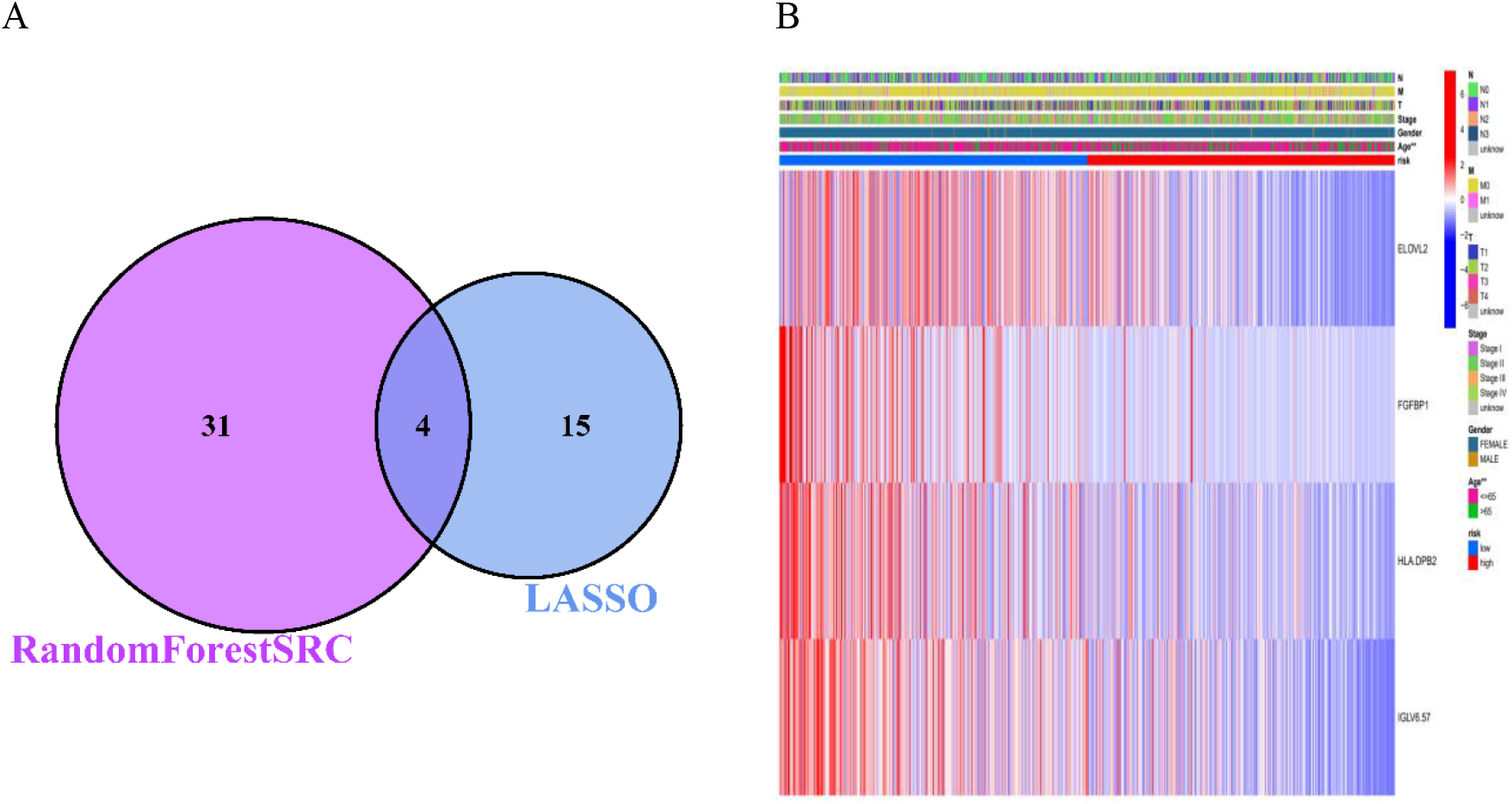
Construction of risk signature in the TCGA cohort. **A** The Random Forest identified 31 genes, and the Lasso regression identified 15 genes. Four genes were obtained from the intersection of the two groups, namely significant prognosis-related genes. **B** Heatmap (Blue indicates lower expression, while red indicates higher expression) for the connections between the clinical features and the risk groups (\**p* < 0.05, \*\**p* < 0.01)

According to the median risk score −1.192541, all patients were assigned to high- and low-risk score groups.

### Evaluation and analysis of the risk model

The GEO cohort with 104 patients Was used as the validation set. Firstly, the *risk score* values of these patients were calculated through the weights trained from the TCGA cohort. Then, the median value of TCGA patients’ risk scores can be regarded as the threshold of low- and high-risk scores. The results indicated that there are 45 cases of low-risk and 59 cases of high-risk in GEO cohort, whereas the TCGA cohort contains 535 and 534 cases of low- and high-risk, respectively. It was found that with the increase of scores, the number of dead patients increased. (Figure 5A-D)

**Figure 5.**
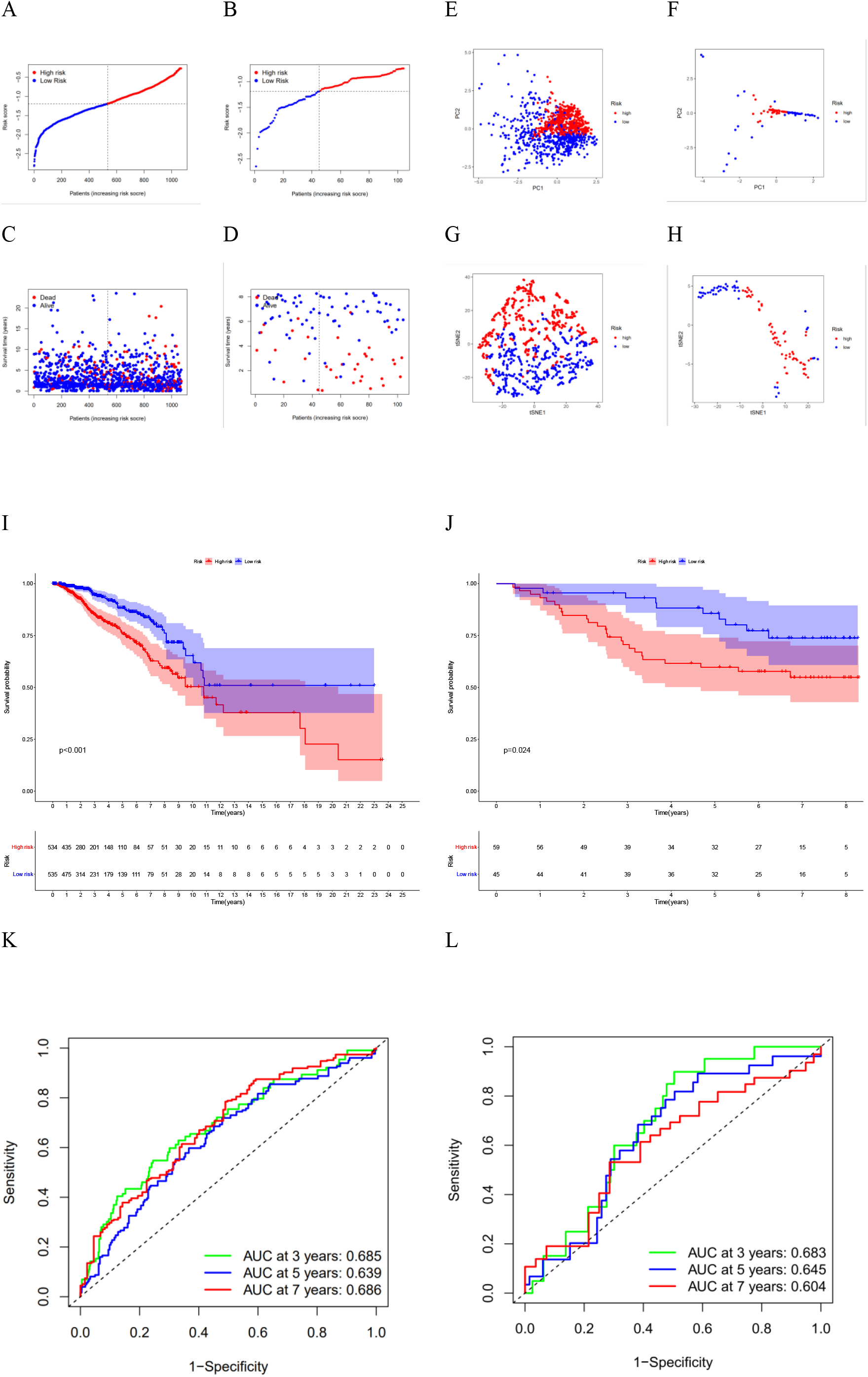
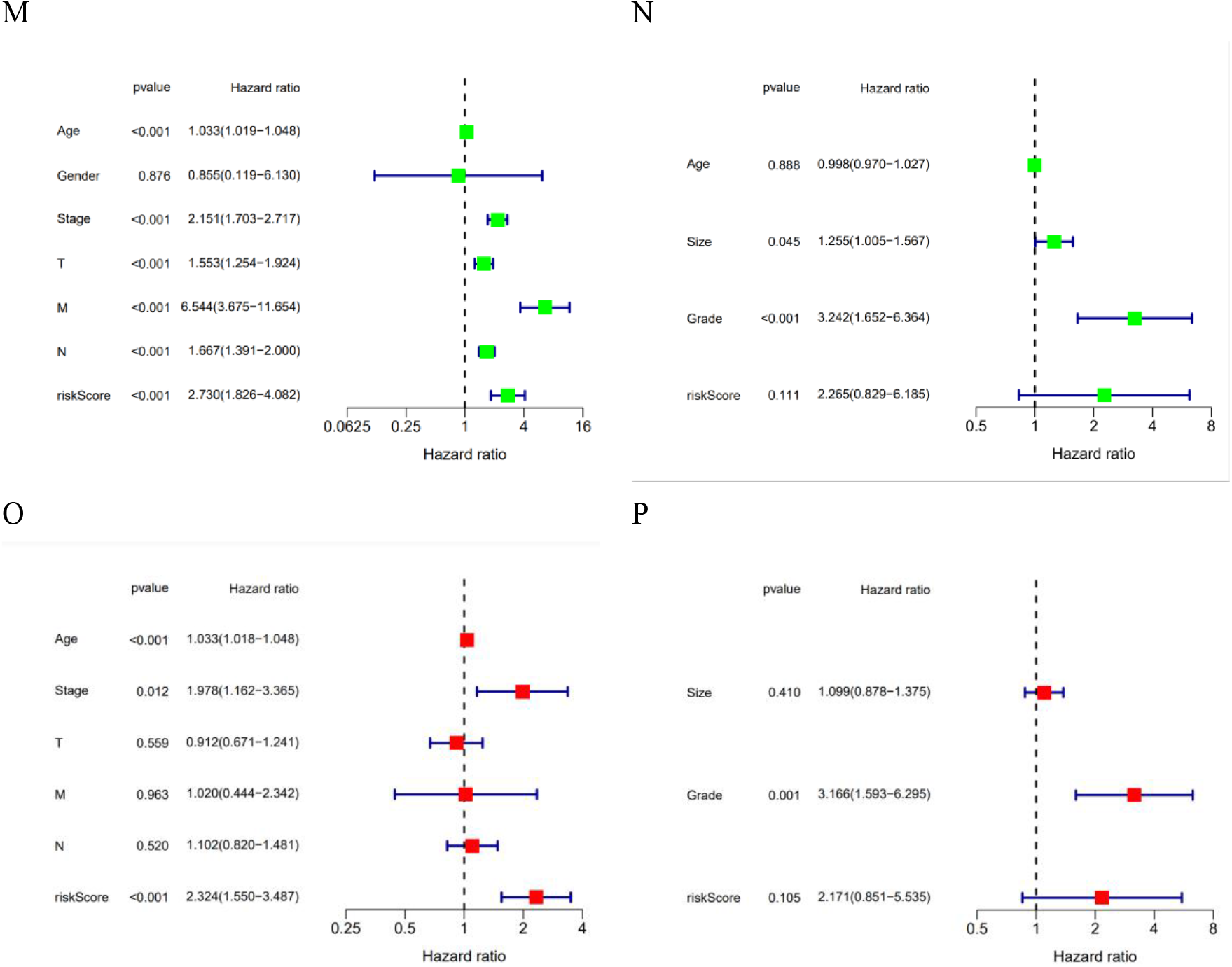
Validation of the risk model, the left is TCGA cohort, and the right is GEO cohort. Patients based on the median risk score(−1.192541) were divided into high- and low-risk groups (low-risk population: on the left of the dotted line; high-risk population: on the right of the dotted line). **A**, **B** Risk curves. **C**, **D** The survival status for each patient (low-risk population:on the left of the dotted line; high-risk population: on the right of the dotted line). **E F** PCA distribution of patients from different sub-groups. **G H** t-SNE distribution of patients from different sub-groups. **I J** Kaplan– Meier curves for the OS of patients in the high- and low-risk groups. **K L** ROC curves for the OS of patients in the high- and low-risk groups. **M N** Univariate COX analysis. **O P** Multivariate COX analysis.

Additionally, the Risk Score takes into account the heterogeneity of patients, the PCA and t-SNE were performed to overview the distribution of patients from different sub-groups, the results showed that patients in different risk sub-groups can be effectively separated based on our prognostic model. (Figure 5E-H)

Moreover, the Kaplan-Meier survival curve was used to evaluate the difference of survival between two different risk sub-groups. According to the curves, the overall survival time of high-risk group is significantly lower than the low-risk group in the both of TCGA and GEO cohorts, (p<0.001 and p<0.021, respectively) which indicate that the effectiveness of 4 final *significant prognosis-related genes* in predicting the BC. (Figure 5I-J)

The ROC curve analysis of GEO and TCGA cohorts demonstrates that our prognostic model has good performance (Figure 5K-L). Furthermore, ROC data were collected from other literatures^[11–13]^ on BC prognostic model of final *significant prognosis-related genes*. After extracting the same data set GSE42568, a horizontal comparison was made. The results show that the mean value of our three-year Time AUC in TCGA and GEO cohorts is 0.67 and 0.644 respectively, which is basically higher than that of the same type of prediction model. (Table 3)

**Table 3.** Comparison results of Time AUC values

| Literature | content | Time AUC Value |  |
| --- | --- | --- | --- |
|  |  | TCGA Cohort | GEO Cohort |
| [11] | Pyroptosis-Related Genes for Predicting Breast Cancer | 3-years: 0.619 | 3-years: 0.690 |
|  |  | 5-years: 0.620 | 5-years: 0.683 |
|  |  | 7-years: 0.645 | 7-years: 0.665 |
| [12] | pyroptosis-related lncRNAs for constructing a prognostic model | 1-years:0.636 | - |
|  |  | 3-years:0.691 |  |
|  |  | 5-years:0.671 |  |
| [13] | Generation of a Pyroptosis-Related Gene Signature for Breast Cancer Prognoses | 0.652 |  |
| our study | Pyroptosis-Related genes for prognosis Genes for Predicting Breast Cancer | 3-years: 0.685 | 3-years: 0.683 |
|  |  | 5-years: 0.639 | 5-years: 0.645 |
|  |  | 7-years: 0.686 | 7-years: 0.604 |

Further regression analysis verified that our model is an independent factor in predicting the overall survival of BC patients. Additionally, In TCGA and GEO shorts, the HR values of univariate and multivariate cox analysis were 2.730, 2.324, 2.265 and 2.171, respectively. (Figure 5M-P)

#### GO, KEGG, GSEA and ssGSEA analysis based on the risk mode

The Mann–Whitney–Wilcoxon Test method was applied with *p* < 0.05 and |logFC| > 1 to find the *subgroup-DEGs* between the high- and the low-score group. Based on these *subgroup-DEGs*, the GO enrichment analysis and KEGG pathway analysis were conducted. In the Biological Progress, these *subgroup-DEGs* were mainly enriched in the positive regulation of cell activation, while Cellular Component analysis shows that these *subgroup-DEGs* were mainly enriched external side of the plasma membrane. Moreover, the Molecular Function analysis illustrated that they were mainly enriched in the antigen-binding. (Figure 6A-D)

**Figure 6.**
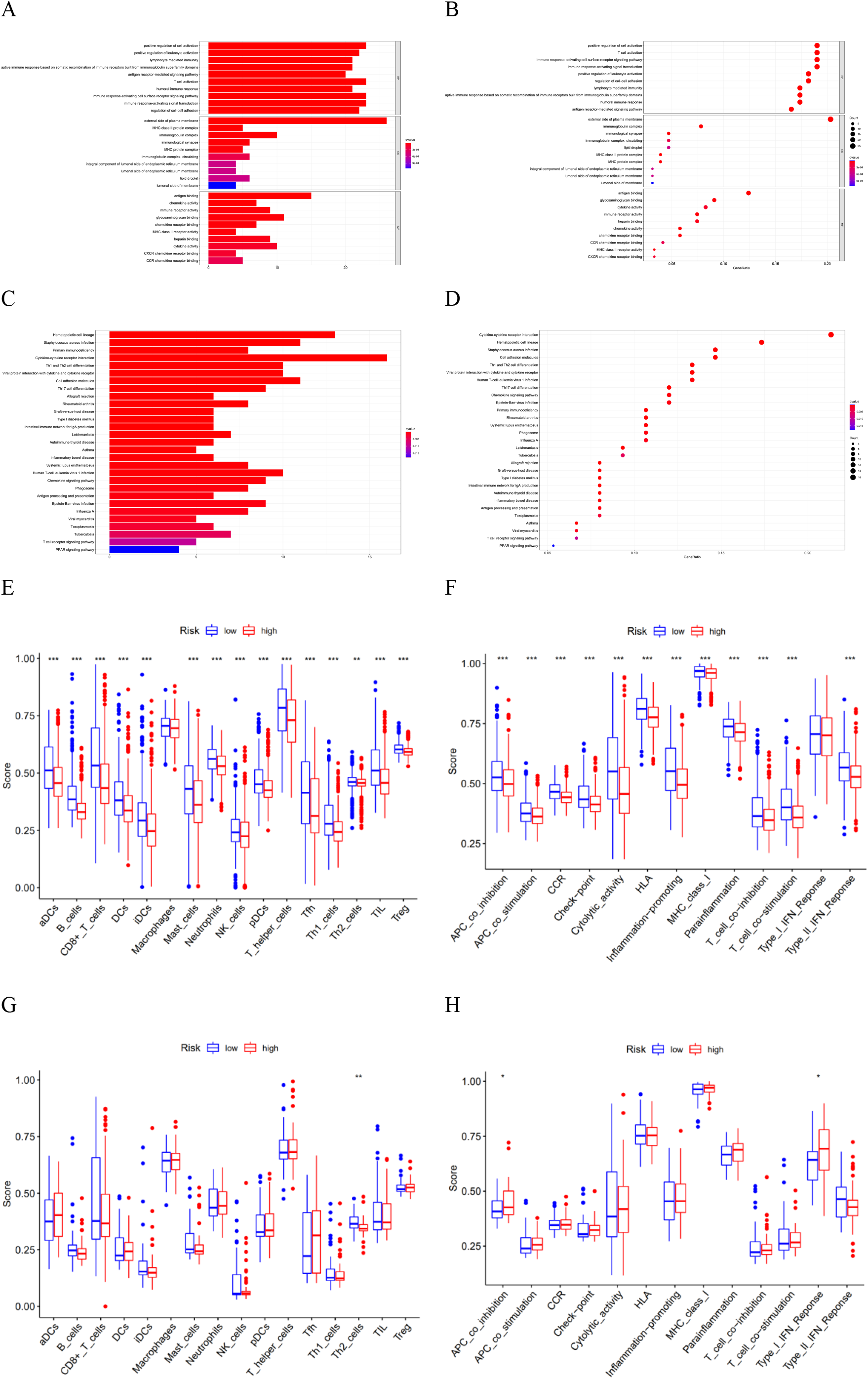
GO, KEGG, GSEA and ssGSEA analysis of subgroup-DEGs. **A, B** GO enrichment analysis and KEGG pathway analysis between low- and high-risk group in the TCGA cohort. **C, D** GO enrichment analysis and KEGG pathway analysis between low- and high-risk group in the GSE42568 cohort. **E, F** Immune microenvironment score analysis of low- and high-risk group in the TCGA cohort. **G, H** Immune microenvironment score analysis of low- and high-risk group in the GSE42568 cohort (\**p* <0.05; \*\**p* < 0.01; \*\*\**p* <0.001).

By performing the ssGSEA, a detailed comparative analysis was made on the enrichment scores of 16 immune cells and activities of 13 immune related pathways of TCGA and GEO cohorts. (Wilcoxon test, \**p* < 0.05; \*\**p* < 0.01; \*\*\**p* < 0.001) The high-risk sub-group generally had lower levels of infiltration of immune cells, especially of aDCs, B cells, CD8+ T cells, DCs, iDCS, Macrophages, Mast cells, Neutrophils, NK cells, pDCs, T helper cells, Tfh, Th1 cells, TIL, Treg than the low-risk subgroup. However, in the GEO cohort, the high-risk sub-group still had high levels of infiltration of immune cells, like Tfh (T follicular helper cells), aDCs (activated DCs) and DCs (dendritic cells), which should be paid attention to. Besides, all the immune pathways showed lower activity in the high-risk group than in the low-risk group in the TCGA cohort. When assessing the immune status in the GEO cohort, some pathways showed relatively opposite sides. For instance, the type-1 IFN response and the APC co inhibition pathways showed higher activity in the high-risk group than in the low-risk group.(Figure 6E-H)

## Discussion

BC is one of the most common cancers in women with a high mortality rate and seriously endangers health. Until now, the treatment of BC still faces enormous challenges. On the one hand, it is necessary to consider the high metastatic heterogeneity of cancer, on the other hand, the research of immunotherapy is still under study. Therefore, it is urgent to model and analyze the prognosis through gene selection, so as to strengthen the understanding of the prognosis of patients.

In this study, a prognostic model based on 4 *significant prognosis-related genes* was proposed to predict the BC prognosis. The TCGA cohort was used as the training set of the model, and a risk score oriented prognostic criterion was proposed. At the same time, the GEO cohort was introduced to verify the effectiveness of the model with different indicators. The results showed that by comparing with other models, our prognostic model can more accurately divide patients into different sub-groups, which has positive clinical significance for subsequent treatment.

In terms of pyroptosis, it is considered to be a programmed cell death, which may be caused by different pathological stimuli. It is characterized by cell expansion and cell membrane rupture. Due to the release of cell fluid, pyroptosis is recognized as a kind of inherent inflammation, which is of great significance to the proliferation and migration of various cancer cells. Besides, many literatures show that it has certain value in the molecules promoting pyroptosis, including non-coding RNA, and has certain significance in cancer treatment^[14–15]^. Current studies have shown that the occurrence of tumor is closely related to the pyroptosis, which has a certain inhibitory effect on the emergence and growth of tumors in a variety of cancers, such as lung cancer, leukemia, melanoma and other tumors^[16–27]^. pyroptosis has a certain expression in the genes of BC patients, and the relevant literature has introduced and information mining^[11–13, 28–30]^. In the study of BC, the over expression of GSDMB is related with poorer prognosis and tumorous progression^[31]^. moreover, the regulation of tumor immune microenvironment promotes inflammasome/IL-1 pathway to play an important role in prgression and invasion of BC^[32]^.

BC prognostic modeling analysis of pyroptosis has a certain research basis in 2021. For instance, Ye Tian et al.^[28]^ found 9 LncRNAs that had a significant effect on overall survival through Cox analysis, which were LMNTD2-AS1, AL589765.4, AC079298.3, U62317.3, LINC02446, AL645608.7, HSD11B1-AS1, AC009119.1, and AC087239.1. These LncRNAs are used to build the risk score model. Shuang Shen et al.^[11]^ extracted four genes CASP9, ELANE, GPX4, and IL18, and constructed them as a prognostic model. By performing LASSO and Cox regression analysis, they divided 1034 patients into different sub-groups. Lili Gao et al.^[29]^ identified seven pyroptosis-related lncRNAs, including AC121761.2, AC027307.2, LINC01871, U73166.1, AL513477.2, AC005034.5 and AL451085.2. They believe that these lncRNAs are closely related with the survival of BC patients. Wenchang Lv et al.^[12]^ built a prognostic model according to 8 pyroptosis-related lncRNAs (AC004585.1, DLGAP1-AS1, TNFRSF14-AS1, AL606834.2, Z68871.1, AC009119.1, LINC01871 and AL136368.1.) they found. Dandan Xu et al.^[13]^ established a three-gene regression model. The IL18, GSDMC, and TIRAP was considered to be an independent risk factor for prognosis of BC.

Generally speaking, our work can be divided into the following parts. First, the complete RNA sequences were obtained from TCGA cohort. These samples contained 39740 different genes. Through data extraction, it was organized into Gene Expression Matrix. In order to explore the influence of *pyroptosis-related genes* in BC, the intersection of *pyroptosis-related genes* and genes was extracted from the Gene Expression Matrix. The results showed that 33 *pyroptosis-related genes* have a certain influence in BC. By Mann – Whitney – Wilcoxon Test (parameter *p* < 0.05), the genes were further extracted into 26 *pyroptosis-related differential genes* related to BC. These genes were analyzed and displayed in more detail, including PPI, correlation network and Heatmap. By univariate Cox regression, 3 *pyroptosis-related prognostic genes* were obtained, which are significantly different between normal and BC patients.

Then, different clustering algorithms were performed to group these BC patients into different sub-types. Through clustering, BC and screened 768 *subtype-DEGs* were further explored through Mann – Whitney – Wilcoxon Test. In the clustering study, not only the Consensus Clustering was performed, but also two other techniques for classification verification. Therefore, the EM and XMeans were conducted on the basis of not specifying the number of sub-types respectively. The results showed that it is best to divide BC patients into two categories, which is also consistent with Consensus Clustering.

Next, the BC prognostic model was constructed for final *significant prognosis-related genes* to distinguish the degree of disease in patients. After extracting relevant features through random forest and lasso regression methods, the following: ELOVL2, IGLV6-57, FGBP1, and HLA-DPB2 were obtained.

Furthermore, the TCGA and GEO cohorts have verified the effect of the model on ROC curve, Survival curve, Risk analysis and PCA demonstration, respectively.

Finally, based on the classification features of the model, the Gene Expression Matrix was extracted from TCGA cohort again, obtained the subgroup-DEGs according to the high- and low-risk score sub-groups, and further discussed and analyzed the BC. Biological function analysis and immune microenvironment analysis were carried out, such as GO and KEGG enrichment analysis and ssGSEA analysis.

Although our study may have a good clinical guiding, there are still several limitations. Firstly, the patients were divided into two sub-types. Although it has been verified that the survival of the two subtypes of disease is significantly different, the essence of the BC disease subtype is still unclear. Secondly, due to the limited clinical characteristics of patients, it was unable to conduct Cox analysis on more factors, especially in the analysis of GEO cohort. Thirdly, it is difficult to ascertain the role of four genes in the pathway of pyroptosis death in BC due to a lack of data.

## Conclusion

With the progress of programmed cell death research, pyroptosis, a method of cell death that plays a dual role in tumor discovery, development, and treatment, has gradually attracted the attention of researchers. In this paper, BC patients were divided into different sub-types. “ELOVL2”, “IGLV6-57”, “FGBP1” and “HLA-DPB2” genes were fully excavated as the independent risk factors of BC diagnoses. At the same time, the corresponding prognostic model was constructed, which can effectively guide the division of patients. In addition, the differences of biological function and immune microenvironment were explored through high- and low-risk grouping.

